# A spiking LIF model captures the role of Somatostatin and Parvalbumin neurons in generating oscillations in V1

**DOI:** 10.1101/2025.01.24.634718

**Authors:** Dario Milea, Nicolò Meneghetti, Alberto Mazzoni, Enrico Cataldo

## Abstract

**Purpose:** Oscillations in the primary visual cortex of the mammalian brain have been demonstrated to arise from the balance between excitatory and inhibitory activity. Experimental studies suggest that different inhibitory neuron populations might make specific contributions to such oscillations, but the underlying mechanism has not yet been assessed.

**Methods:** We modified a standard excitatory-inhibitory spiking neuron model of layer 4 of the primary visual cortex and we investigated the effects on oscillations of the differentiation of inhibitory neurons in somatostatin and parvalbumin neurons.

**Results:** Our model reproduced the hypothesis that somatostatin and parvalbumin neurons are responsible for beta (15-25)Hz and gamma (40-70)Hz band oscillations, respectively.

**Conclusion:** To date, this is the simplest model accounting for this phenomenon and could therefore be suited to study pathologies in which the two populations have specific roles, such as migraine.

## 1 Introduction

Neural oscillations are routinely observed in various brain regions, including cortical and sub-cortical areas (Buzsaki, 2019, 2006; Buzsaki and Draguhn, 2004; Buzsaki et al., 2012; Jadi and Sejnowski, 2014), and the mammalian primary visual cortex (V1) exhibits a rich repertoire of distinct oscillatory phenomena (Mazzoni et al., 2011).

Many studies have investigated the deep correlation between characteristic neuronal oscillations and sensory, motor or cognitive functions (Gray and Singer, 1989; Sanes and Donoghue, 1993; Fries et al., 2001). For instance, Kayser et al. (2012); Montemurro et al. (2008) suggest that oscillatory rhythm in a low frequency range (*<* 12 Hz) is the main platform for information coding, whereas the gamma frequency band has a strong correlation with visual attention (Engel et al., 2001). Recent studies (Meneghetti et al., 2021; Veit et al., 2017, 2023; Domhof and Tiesinga, 2021) show how low and high gamma frequency bands can be considered distinct but complementary means to process visual stimuli.

In different mammalian species, such as mice (Meneghetti et al., 2022) and monkeys, neurological disorders appear to be correlated with altered neuronal activity. These phenomena have also been observed in humans, such as Parkinson’s disease and migraine (Liguz-Lecznar et al., 2022; Marchionni et al., 2022; Vreysen et al., 2016). However, the principles that regulate the rhythmic activity of neural populations are still debated.

In recent decades, studies by Brunel and Wang (2003); Brunel (2000); Brunel and Sergi (1998); Renart et al. (2019) modeled a network of spiking neurons that were able to reproduce the main cortical oscillations. The system was composed of two neural populations, one excitatory and the other inhibitory, connected to each other with a homogeneous probability. A Poisson process represented the main excitatory input from a wide external world that started and maintained, under certain circumstances, the network activity. Despite the use of just two distinct populations, the system revealed a rich repertoire of different behaviors, and the use of the leaky-integrate-and-fire point-neuron model made the system simplified enough for an analytical study. However, since the model was used on only one inhibitory population, the network lacked differentiation among distinct rhythmic activities.

Recently, Billeh et al. (2020) developed a model of a portion of the mouse primary visual cortex. The model incorporates all six layers of the visual cortex and is stimulated by a module that mimics the lateral geniculate nucleus. The model is one of the most accurate models ever used to reproduce primary visual cortex oscillations, but it is also computationally expensive and requires various tests to fit the parameter space with data. The use of a detailed neural model results in a large parameter space, making the system difficult to control and explore. Furthermore, the model fails to reproduce one of the most fundamental features of the real visual cortex: neural population gamma rhythms.

Recent years have seen significant efforts to understand the mechanisms by which different inhibitory interneurons, when stimulated in different layers of the visual cortex, influence cortical oscillations (Wagatsuma et al., 2022; Hahn et al., 2022).

In the present study, we developed a network model of layer 4 of the mouse primary visual cortex. This model comprised one class of excitatory neurons and two distinct inhibitory neuron types: parvalbumin and somatostatin neurons (Lee et al., 2010; Tremblay et al., 2016; Song et al., 2020; Lukomska et al., 2020).

In mouse V1, these neuron types constitute the three major classes, exhibiting significant differences in their spiking activities, synaptic connectivity, and functional roles (Miao et al., 2016; Pfeffer et al., 2013; Cottam et al., 2013). Parameter values were obtained from the most up-to-date database (Campagnola et al., 2022). Previous studies by Chen et al. (2017) and Veit et al. (2017) suggest that these interacting neuronal populations play a crucial role in regulating synchronized oscillatory activity when an input stimulus falls within their receptive fields.

The use of a simplified single-neuron model and a randomly interconnected network has allowed us to perform analytical studies of the system in addition to simulations. Our results suggest that somatostatin and parvalbumin neurons contribute primarily to beta (15-25)Hz and gamma (40-70)Hz band oscillations, respectively. These findings highlight the crucial role of distinct inhibitory neuron subtypes in modulating the rhythmic activity of the cortex.

## 2 The Model

### 2.1 Single Neuron

We describe single neurons using the Leaky Integrate-and-Fire (LIF) model. The LIF model provides a good balance between biological realism and computational efficiency (Izhikevich, 2004; Burkitt, 2006; Teeter et al., 2018). Each neuron in our model must satisfy the following membrane potential equations:

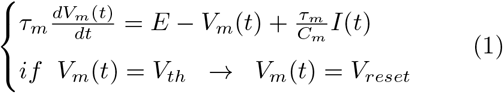

where *τ*_*m*_, *C*_*m*_ and *E* represent the membrane time constant, capacitance, and resting potential, respectively. *V*_*th*_ denotes the membrane threshold, *V*_*reset*_, the membrane reset value, and *δt*_*ref*_, the refractory time that silences the membrane after a spike. Every time the membrane potential reaches the threshold, a spike is emitted, and *V*_*m*_(*t*) is immediately reset to *V*_*reset*_. The neuron is inhibited and does not react to other inputs for a duration of *δt*_*ref*_. The term *I*(*t*) represents a generic current stimulus, which will be further elaborated upon in the next section.

Parameter values for parvalbumin, somatostatin and excitatory neurons are calculated as a weighted average of the neural subclasses within each population, according to Campagnola et al. (2022) (see table 1).

**Table 1.**
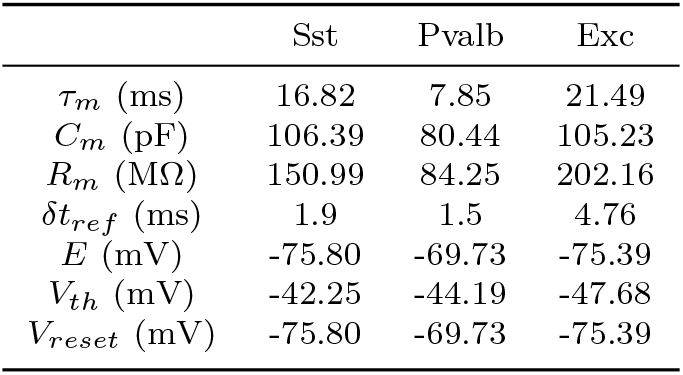
Neural parameter values. From left to right, we show the somatostatin, parvalbumin and excitatory neural parameters. The rows represent the membrane time, capacitance and resistance, refractory period, resting potential, threshold and reset potential according to Campagnola et al. (2022).

As a first step, we characterize the dynamics of the single neural model, as described in Appendix A.

### 2.2 Synapses

In the present model, the term *I*(*t*) represents a synaptic current: *I*(*t*) = *I*_*syn*_(*t*), that includes all synaptic influences. These are the currents induced by the interaction with the other neurons of the network and the current mediated by the external synapses that are supposed to be elicited by the Lateral Geniculate Nucleus in our model. Then, the synaptic current can be written in the form:

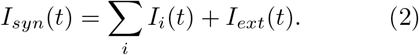

The mathematical form of each synaptic term on the r.h.s. of Eq. (2) is the following:

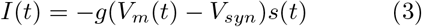

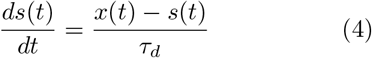

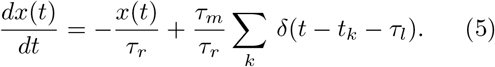

The parameter *g* represents the synaptic conductance, which determines the strength of each individual synaptic connection. Synaptic kinetics is characterized by two parameters: the rise time *τ*_*r*_ and the decay time *τ*_*d*_. The latency *τ*_*l*_ represents the delay between spike emission and arrival at the target neuron. The time of the *k*-th spike is denoted by *t*_*k*_. *V*_*syn*_ stands for the reversal potential of the synaptic channel, which is 0 *mV* (− 70 *mV*) if the synapse is excitatory (inhibitory) (Arkhipov et al., 2018). The final term in equation 5 represents the external excitatory input, modeled as a Poissonian process. This input simulates the thalamic drive to the primary visual cortex during high-contrast visual stimulation, as described by Meneghetti et al. (2021, 2022). Finally, *x*(*t*) and *s*(*t*) are two functions related to the time course of the synaptic current. The signs of the currents follow the electrophysiology convention.

Table 2 shows the parameter values for external synapses (Kloc and Maffei, 2014; Billeh et al., 2020). Note that the somatostatin neurons do not receive any kind of excitation from other brain regions in our model. Moreover, no latency is specified for the external synapses, simply being a delay constant of the network response to the stimulus.

**Table 2.**
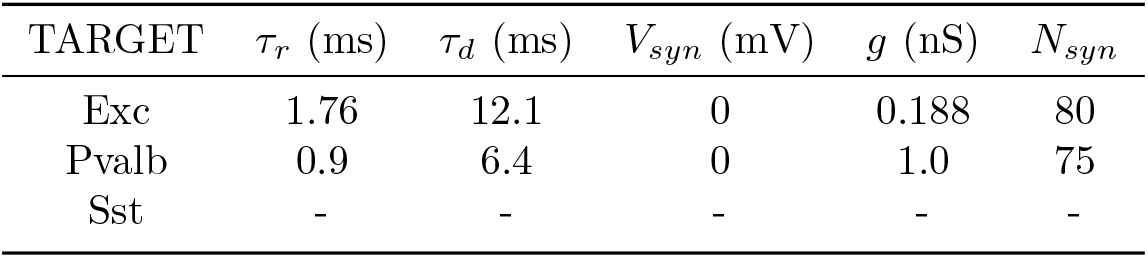
Parameter values of external synapses. From left to right are shown the rise time, the decay time, the reversing potential, the conductance and the number of synapses per neuron. The neural classes refer to the target neuron to which the synapses are connected. The characteristic times are acquired from Kloc and Maffei (2014), the resting potential and the number of synapses are acquired from Billeh et al. (2020). We deduce the conductance to obtain a mean firing rate per neuron comparable to that registered in Billeh et al. (2020).

| TARGET | $\tau_r$ (ms) | $\tau_d$ (ms) | $V_{syn}$ (mV) | $g$ (nS) | $N_{syn}$ |
| --- | --- | --- | --- | --- | --- |
| Exc | 1.76 | 12.1 | 0 | 0.188 | 80 |
| Pvalb | 0.9 | 6.4 | 0 | 1.0 | 75 |
| Sst | - | - | - | - | - |

Synaptic parameter values for intra-network connections are summarized in figure 1. Decay times, rise times and latencies are shown with reference to Campagnola et al. (2022). We call *V*_*target*_ the parameter that stands for the maximum value of the Post Synaptic Potential. It captures the strength of a synaptic response to a presynaptic spike. It is measured with respect to the equilibrium membrane potential, which we set equal to 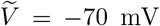 (− 55 mV) for the excitatory (inhibitory) synapses, respectively, as in Campagnola et al. (2022). Synaptic conductances are determined by adjusting *V*_*target*_ to achieve the desired PSP amplitude following a single presynaptic spike, as outlined in the Methods section of Billeh et al. (2020). For consistency checks, simulations are performed using the exact expressions for the synaptic currents. Unlike external inputs, intra-network connectivity is defined probabilistically. Instead of specifying the number of synapses, we use a connection probability matrix. For example, if the populations 𝒜 and ℬ are interconnected and the number of neurons of the populations is *N*_*𝒜*_ and *N*_*ℬ*_, respectively, then the number of connections that every neuron of *ℬ* receives from *𝒜* is on average *N*_*𝒜*_*𝒫*_*𝒜ℬ*_, while the number of connections that every neuron of *𝒜* receives from ℬ is on average *N*_*ℬ*_ *𝒫*_*ℬ𝒜*_, and the probability of connection is in general asymmetric.

**Fig. 1.**
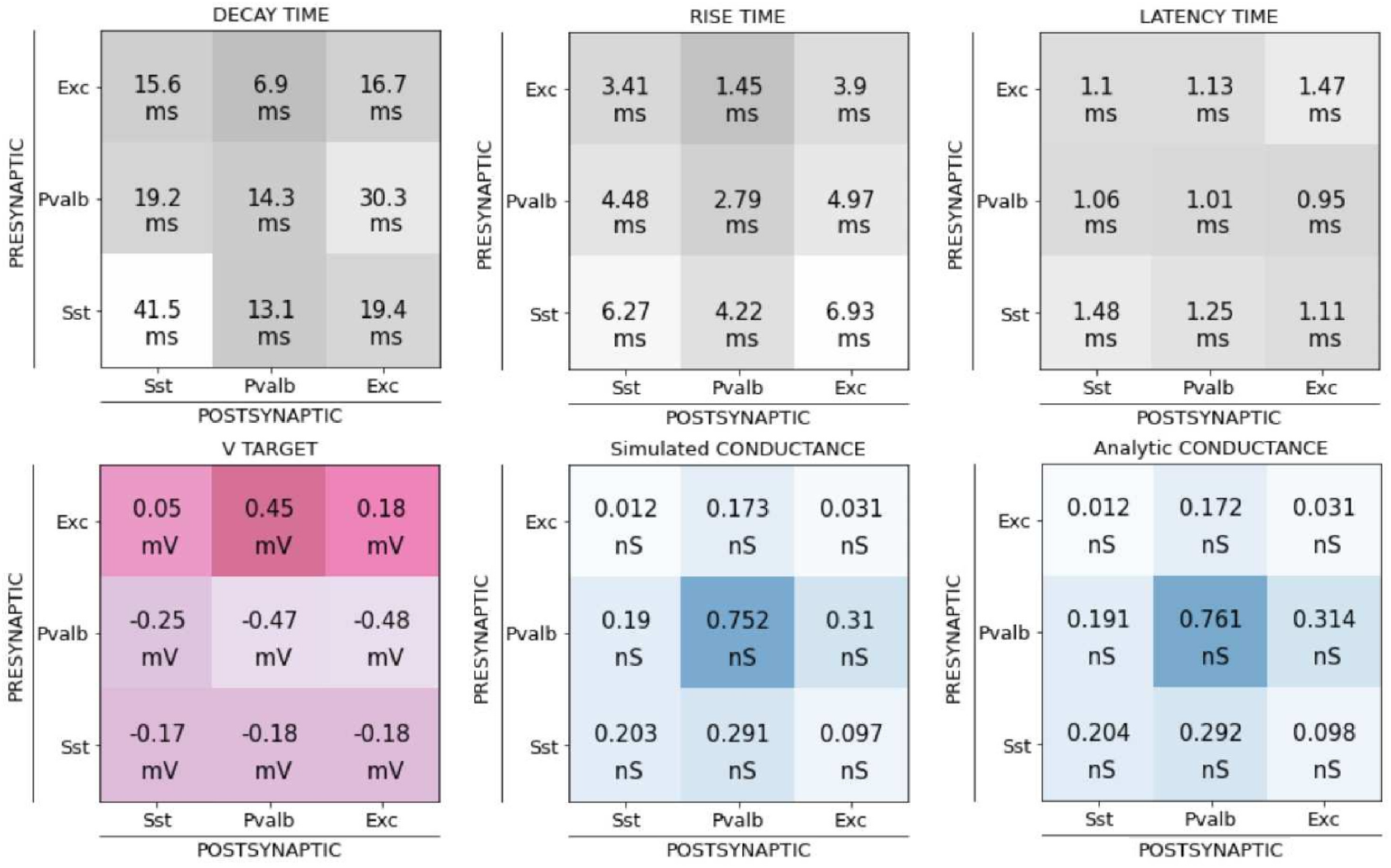
Parameter values of the synapses among neurons. The kinetics parameter values of the new model are shown in the first row. From left to right the decay time, the rise time and the latency - note the presence of the latency time that we have not used in the external synapses instead -.The second row displays the target value of the membrane potential after a spike is coming. All these values are taken from Campagnola et al. (2022). In blue shades, we report the conductances, derived analytically and by simulations. We enter on the left of each matrix with the presynaptic neuron and go down to the postsynaptic neuron.

### 2.3 The network

When neurons are interconnected, the equations from 1 to 5 remain valid, but the sum over the spike train makes it prohibitive to find an analytical solution. However, under specific parameter regimes, the population activity of the network exhibits oscillatory behavior.

Population activity represents the instantaneous firing rate of the entire network. The recurrent connections make the output of one neuron to serve as input to the others, and drive the network into a state of rhythmic activity. We can stress this behavior by writing the current equation 5 as Brunel and Wang (2003) suggest:

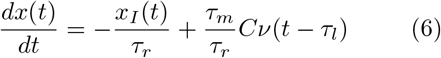

where we approximate that the system response has a perfect sinusoidal form at frequency *ω*:

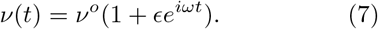

The variable *C* = 𝒫*N*, where 𝒫 is the connection probability and *N* is the number of neurons for a single class, represents the average number of connections for each neuron, and in general it depends on both the presynaptic and postsynaptic neuronal class. For this reason, to be as general as possible, we specify both the firing population and the target population with two capital letters as subscripts, where necessary.

The target model is composed of 10000 cells, 85 % excitatory, 10 % parvalbumin and 5 % somatostatin (Lee et al., 2010; Schüz and Palm, 1989). From now on, we refer to these neural classes with the letters *E, P* and *S*, respectively. The external excitatory input is usually set as a Poissonian input with a mean of 7.3 Hz as suggested in Schultz et al. (2016). Because the connectivity is not yet completely known and there are many different models in the literature (Brunel and Wang, 2003; Billeh et al., 2020), we change 𝒫 to fit the biological data from Billeh et al. (2020).

To address the complexity of the final network of three distinct populations one step at a time, we first study the case of a two-population network.

## 3 Results

### 3.1 Two populations

We used parvalbumin neurons as the sole inhibitory cell type in our initial simulations. The number of parvalbumin neurons (1000) and excitatory neurons (8500) was fixed at their target values. This resulted in a well-defined balance between excitation and inhibition from parvalbumin cells to excitatory neurons. This established baseline activity will not be significantly altered when the third neural population, somatostatin neurons, is introduced. Both populations received the same external input, modeled by a Poisson firing rate with a constant mean.

We begin our analysis by studying the network dynamics at increasing levels of complexity. First, we analyze the interactions within the basic excitatory-inhibitory loop. Next, we incorporate recurrent inhibition into the model. Finally, we include recurrent excitation to complete the network connectivity. The main equations governing the dynamics of the network at each level of connectivity are derived in B.

Figure 2 illustrates the analysis results on the network’s oscillatory behavior. The analysis focuses on the phase lag between input stimulation and the network’s response to predict the peak oscillation frequency.

**Fig. 2.**
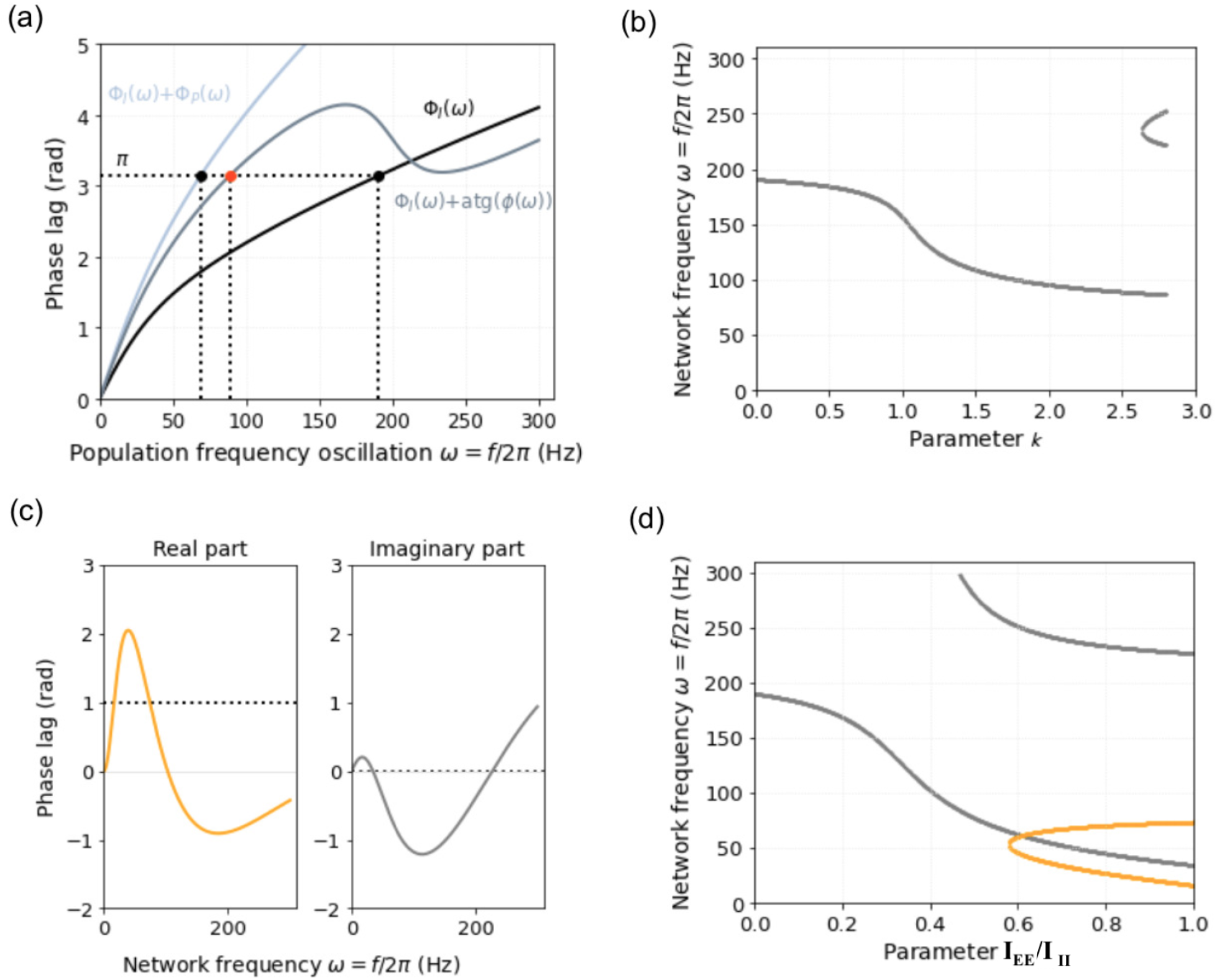
Phase lag analysis for different networks composed of two populations. (a) Solutions of the equations describing the phase lag in three different cases. The black continuous line shows the Φ (*w*) = *π* function in the case of a completely inhibitory network. It stands for the phase lag from the presynaptic input and the postsynaptic output of the network oscillation. The light-gray is the phase due to the inhibitory-excitatory cell loop, without any recurrent activity (see Eq. B11). Finally, the dark-gray line represents the case of an inhibitory-excitatory loop with recurrent inhibition for *k* = 2, as shown in Eq. B17. The intersections between the phase lag curves and *π*, highlighted as a black dashed line, display the frequency population oscillation we expect to find. (b) The case of an inhibitory-excitatory loop with recurrent inhibition has a phase lag that depends on the parameter *k*. Here, the behavior of the population frequency as a function of this parameter is outlined. (c) The fourth case refers to complete connectivity; see Eq. B23 in the Appendix. The real and imaginary parts of the phase lag function are shown for *A* = 5 and *I*_*EE*_*/I*_*II*_ = 1. The population frequency is computed by equalling the real part to the value of one and the imaginary part to zero. (d) The solutions are shown for *A* = 5 in the respective colors as a function of the parameter *I*_*EE*_ */I*_*II*_. Note that just for one value, the real and imaginary parts share the same network frequency, which is the one we expect to find by simulations.

For a purely inhibitory network (black line in figure 2(a)), a high oscillation frequency of around 185 Hz is expected. When the excitatory-inhibitory loop is studied the frequency is reduced to 70 Hz, (light-gray line). Adding recurrent inhibition to the loop leads to a compromise frequency of around 85 Hz (red dot in figure 2(a)), which depends on a parameter *k*, see equation B18. Interestingly, for stronger values of k, the expected oscillation frequency shows a branching behavior (figure 2(b))

The fully connected network’s frequency depends on the ratio of excitatory and inhibitory currents (*I*_*EE*_*/I*_*II*_) and an amplification factor (*A*). Figure 2(c) shows both the real and imaginary parts of the phase lag for a specific amplification (*A* = 5) and a current ratio of *I*_*EE*_*/I*_*II*_ = 1. In figure 2(d), with a constant amplification factor, the network frequency is expected to be around 60 Hz when the current ratio increases from zero to 0.62 (*I*_*EE*_*/I*_*II*_ *≈* 0.62).

To investigate the effects of the new neuron and synapse types, we utilized a fully connected network as our primary platform. As detailed in Appendix B.4, a systematic study led to the following connection probabilities: *𝒫*_*P,P*_ = 0.4, 𝒫_*E,P*_ = 0.42, 𝒫_*P,E*_ = 0.15, 𝒫_*E,E*_ = 0.22.

Figure 3 summarizes the key results for this two-population network. Panel (a) illustrates the network connectivity. Panel (b) presents the corresponding raster plot, generated by stimulating the network with an external Poissonian input at a mean firing rate of 7.3 *Hz*. Panel (c) depicts the activity of individual neurons, revealing lower activity levels in excitatory neurons compared to parvalbumin neurons. The power spectrum in panel (d) reveals a main peak at approximately 5 Hz. Additionally, the spectrum exhibits other prominent oscillations in the gamma range, around 60 Hz. When we decrease the external stimulation to 5.5 Hz, the power spectrum becomes flat, and the network activity significantly diminishes. This low-activity state, characterized by minimal network activity, serves as the baseline for subsequent analyses of network dynamics.

**Fig. 3.**
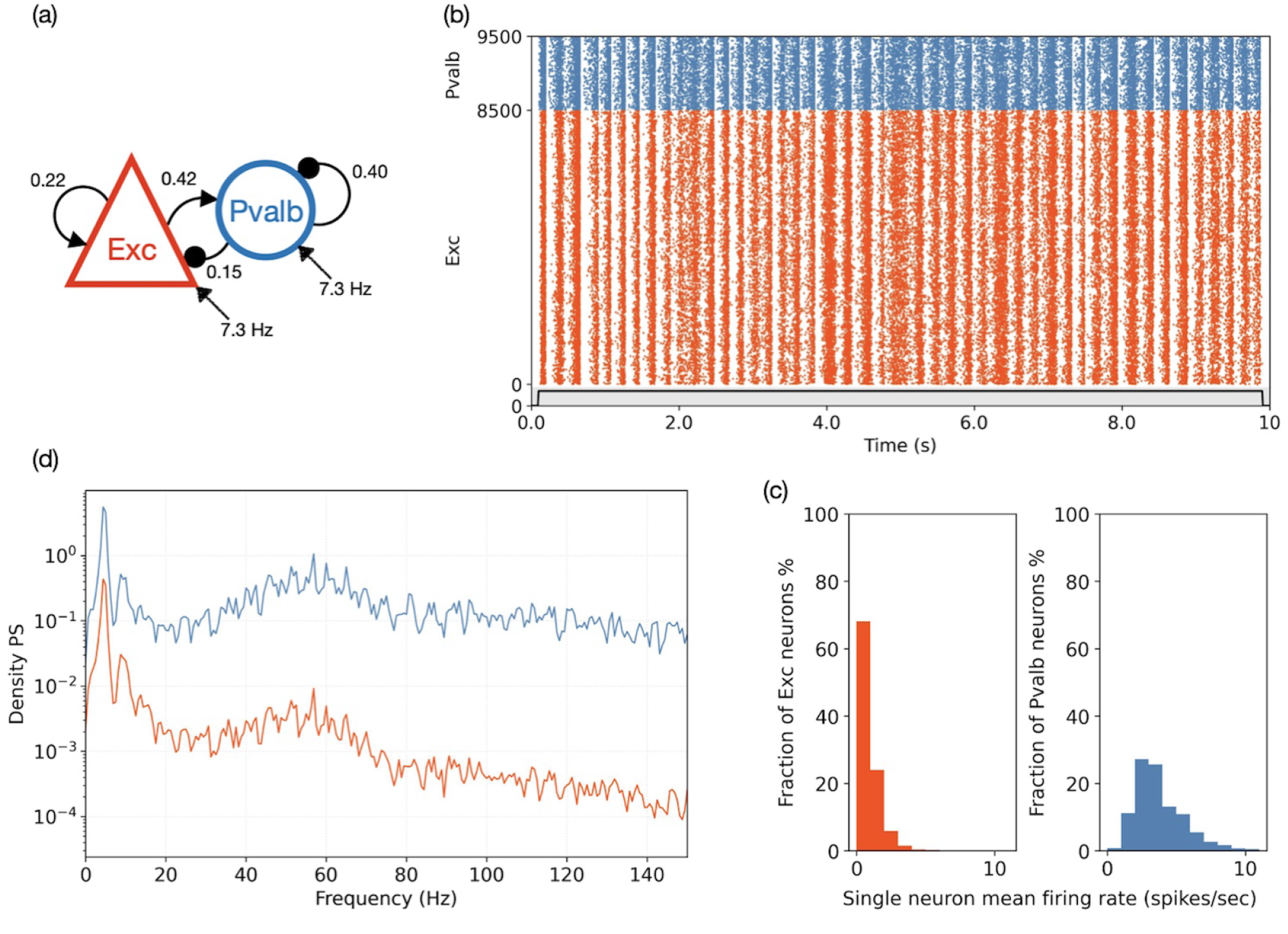
Pvalbumin-excitatory full connectivity model. (a) Schematic of the heterogeneous connection probability. The arrows from below represent for the external synapses, 75 for the excitatory neurons and 80 for the inhibitory neurons. The mean frequency of the Poissonian stimulus is specified. (b) Raster plot of the entire network made of 1000 inhibitory neurons and 8500 excitatory neurons. The black line at the bottom corresponds to the activation of the external stimulus. (c) Histograms of the single neuron mean firing rate for the excitatory and parvalbumin neurons respectively. (d) Power spectrum (Welch’s method applied) of the population activity. Red indicates excitatory neurons, and blue indictes parvalbumin neurons.

### 3.2 Three populations

The final network comprises 8500 excitatory neurons, 1000 parvalbumin neurons, and 500 somatostatin neurons, representing the two inhibitory classes. The synaptic connectivity within the network is defined in Appendix C. Figure 4 summarizes the key results obtained with this three-population network. Panel (a) illustrates the connection probabilities among the different neuronal populations and the mean frequency of the external Poissonian input. Panel (b) presents the corresponding raster plot. A general feature observed in panel (c) is a decrease in the mean firing rate of single neurons compared to the two-population case, likely due to the increased inhibitory influence of the somatostatin interneurons.

**Fig. 4.**
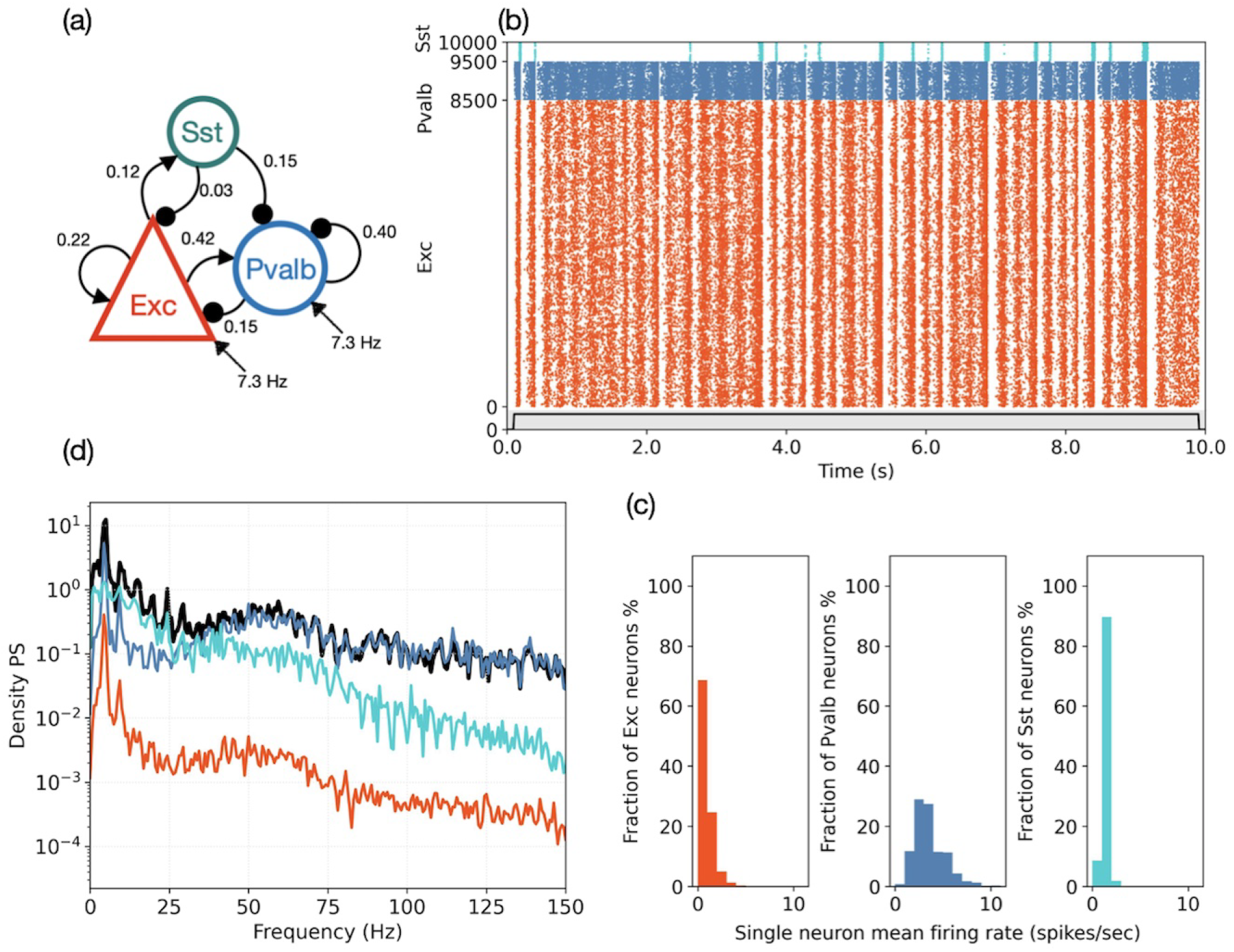
Three-populations model. (a) Schematic of the heterogeneous connection probability. The arrows from below represent the external synapses, 75 for the excitatory neurons and 80 for the inhibitory neurons. The mean frequency of the Poissonian stimulus is specified. (b) Raster plot of the entire network made of 1000 parvalbumin neurons, 500 somatostatin neurons and 8500 excitatory neurons. The black line at the bottom corresponds to the activation of the external stimulus. (c) Histograms of the single neuron mean firing rate for excitatory, parvalbumin and somatostatin neurons. (d) Power spectrum (Welch’s method applied) of the population activity. Red refers to the excitatory neurons, blue to the parvalbumin neurons and gray to the somatostatin neurons. The black line shows the oscillation components of the entire network response.

Several parameters significantly influence single neuron responses. Notably, the connection probability between somatostatin (Sst) and parvalbumin (Pvalb) neurons (*𝒫*_*S,P*_) plays a crucial role. Increasing 𝒫_*S,P*_ generally leads to a gradual increase in the mean firing rate of all neuron types, particularly when Sst-to-Exc inhibition is weak.

However, high 𝒫_*S,P*_ values can induce a sudden and dramatic shift in network activity, a phenomenon similar to a phase transition. (The conductance (g) of synapses from Sst to Exc neurons exhibits a similar effect.) Initially, increasing recurrent inhibition from Sst to Pvalb enhances the activity of Sst and Pvalb neurons. However, stronger Pvalb inhibition reduces the firing rate of excitatory (Exc) neurons, subsequently decreasing Pvalb activity. This dynamic interplay leads to an equilibrium state. This process becomes very intense when 𝒫_*S,P*_ approaches the 0.25 value. A sharp increase in Exc neural activity occurs, Sst neural activity follows, and Pvalb neural activity slows down. Clearly, there is poor inhibition of the Exc neurons to maintain equilibrium, and the inhibition from Sst on Pvalb is strong enough to almost silence the neurons. The inhibition from Pvalb on Exc is then reduced, and the loop Sst - Exc becomes stronger, searching for another state of equilibrium. This phenomenon can be mitigated by increasing recurrent inhibition from Sst to Exc neurons. Stronger Sst-to-Exc feedback effectively counteracts the destabilization, allowing the network to maintain a stable equilibrium.

We can partially mitigate both the increase in neuronal activity and the phase transition by adjusting the excitation of Sst neurons from Exc neurons. If we decrease 𝒫_*E,S*_ Sst neural activity becomes weaker and the phase transition shifts to higher values of 𝒫_*S,P*_, as describe in the supplementary material.

Conductivity, however, appears to have a more pronounced effect when the connection probability *𝒫*_*S,E*_ is stronger.

Focusing on the network activity, see figure 4 panel (d), we observe that Sst neuron activity is primarily associated with beta-band oscillations (15-25 Hz), while Pvalb neuron activity primarily contributes to gamma-band oscillations (40-60 Hz).

## 4 Conclusion

We have described a novel computational spiking neuron network model of layer 4 of the primary visual cortex in rodents composed of three different populations of neurons. Despite its minimalist design, the model successfully captures the relationship between neurophysiological features at the level of single neurons and network behavior and emergent network behavior.

First, we defined the neurons composing the three populations based on groudbreaking neuro-physiological data. Subsequently, we studied their individual and collective behaviors within the network.

When analyzing the neural response function to a Poisson stimulus with a constant mean firing rate (*ν*), we made two approximations (see Appendix A). First, we approximated the Poisson firing rate with a regular firing rate at the same frequency (*ν*). This approximation, valid in the high-frequency regime ((50 − 800) spikes*/*sec), allowed us to derive a simple analytical solution for the neural response function. Second, for low firing rates ((0 − 20) spikes*/*sec), we approximated the Poisson firing rate with white noise. This approximation provided a good fit to the observed data (see Figure S10 in the Supplementary Material).

We began by analyzing a two-population network, incorporating a novel synaptic connectivity scheme between parvalbumin and excitatory neurons. With heterogeneous connectivity, a clear gamma band rhythm in the activity of both populations was visible. Subsequently, we introduced the somatostatin population into the network. Following the establishment of appropriate connectivity among all three populations, the network maintained gamma-band oscillations. Notably, the power spectrum analysis revealed a distinct contribution from the somatostatin population, primarily within the beta-band (below 30 *Hz*). Excitatory and parvalbumin neurons were the primary drivers of gamma-band oscillations, and exhibit a peak around 5 *Hz*.

These findings, demonstrating distinct frequency contributions from different neuronal populations, are consistent with previous experimental studies Chen et al. (2017); Buzsaki (2002); Stark et al. (2013); Cardin et al. (2009); Sohal et al. (2009).

The two different interneuron types were reported to shape the LFPs spectrum differently within the high visual contrast range. Conversely, their specific contribution to the LFPs spectrum in the low contrast range, dominated by the entrainment of gamma oscillating thalamic afferents, was not reported (Saleem et al., 2017; Meneghetti et al., 2021, 2022; Shin et al., 2023). Consequently, we decided not to include the oscillating thalamic inputs at 60 Hz but focused our attention on simulating the responses of our cortical model to increasing levels of Poissonian thalamic afferents, intended to simulate inputs within the high visual contrast range.

We analyzed the way our network responded to the increasing values of the external rate (range 5Hz - 10Hz) as shown in Fig. S1-S2-S3 of supplementary material (Online Resource). We focus on the peak of the oscillation frequency of the populations activity, the amplitude of the oscillation, and the mean firing rate of single neurons. Fig. S2-S3 display the raster plot and its power spectrum in the two limits of the frequency range. Fig. S1 shows how higher values of the external input rate increase both the oscillation frequency of the populations activity and the amplitude of the oscillation, replicating the response captured by Chen et al. (2017).

Recent studies, including those by Chen et al. (2017); Veit et al. (2017), have reported beta-range oscillations in a minority of the recorded excitatory cells and parvalbumin neurons. This trend was not replicated by our model, indicating that the nuanced cellular behaviors in V1 might not be fully captured by our model designed for the overarching trends of the network.

In Chen et al. (2017); Veit et al. (2017) the authors also studied their network behavior when one cell type (parvalbumin or somatostatin neurons) was weakened using optogenetic techniques. To compare our network response with their findings we have added to parvalbumin neurons, or somatostatin neurons respectively, an hyperpolarizing external current, see Fig. S4-S5-S6-S7 in the supplementary material (Online Resource). When the current affected parvalbumin neurons our network showed the characteristic behavior of an oscillator (its main oscillation frequency was approximately 10 Hz) and parvalbumin activity was not reduced. This trend did not replicate Chen et al. (2017); Veit et al. (2017) representing a limitation of our model. However, when somatostatin neurons were hyperpolarized, their mean single neuron firing rate decreased and their activity was progressively silenced. This result is in line with Chen et al. (2017).

In conclusion, our model presents a more biophysically accurate representation of cortical dynamics compared to the previous two-population networks (Brunel and Wang, 2003). In particular, it incorporates three distinct neuronal populations (Chen et al., 2017; Veit et al., 2017; Jang et al., 2020), which align more closely with neurophysiological observations. Although our model is less complex than those developed by Billeh et al. (2020), it exhibits a crucial feature: the emergence of gamma-band oscillations without requiring explicit external gamma-band input. This finding highlights the intrinsic capacity of the network to generate these oscillations through the interplay of its constituent neuronal populations. Furthermore, the model provides a valuable platform for investigating the underlying mechanisms that govern cortical function. Its controllable parameters and simulated behaviors offer a potential avenue to explore the pathophysiology of clinical conditions, particularly migraine. As we have mentioned in the Introduction, the use of parvalbumin and somatostatin neurons as two distinct inhibitory neuronal populations plays a fundamental role in the model of migraine (Liguz-Lecznar et al., 2022; Marchionni et al., 2022; Pietrobon and Moskowitz, 2013; Vreysen et al., 2016). These complex brain disorders are characterized by global dysfunctions in information processing, which can be correlated with the lack of balance in excitatory-inhibitory equilibrium. The underlying cellular mechanism involves both parvalbumin and somatostatin neurons because it alters the regular rhythm of the brain in the gamma and beta ranges, which are generated by the aforementioned neural classes.

## Declarations

### Authors’ Contributions

D.M. wrote the manuscript and prepared the figures. All authors reviewed the manuscript.

## Appendix A Neural model

To characterize the single neuron behavior we use a constant current stimulus of variable amplitude at each trial or a Poisson process with different values of the mean frequency rate (Campagnola et al., 2022).

In the first case, an analytical solution is available; see Eq. A1. The second case is more cumbersome, and we address it using two approximations. When the neuron fires at a rate higher than approximately 50 Hz, we use a constant-input firing-rate approximation equal to the *ν* value and neglect the probabilistic behavior of the external synapses. We are able to find the mathematical expression of Eq. A2.

At the opposite limit we pose two conditions: each incoming spike has little effect on the system; the number of incoming spikes is high in the interval *τ*_*m*_ because the number of external synapses per neuron. Then the membrane evolution can be described as Brownian motion using the diffusion model and the Fokker-Planck equation, as suggested by Brunel (2000); Brunel and Sergi (1998); Gerstner et al. (2014).

Eq. A3 displays the mean first passage time that the neuron takes from the reset potential to the threshold. The integration limits are 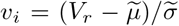 and 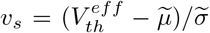, respectively.

Two opposite limits are compared in Fig. S10 in the supplementary material (Online Resource).

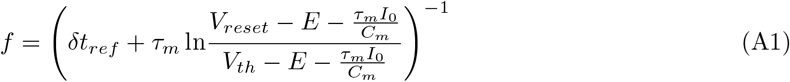

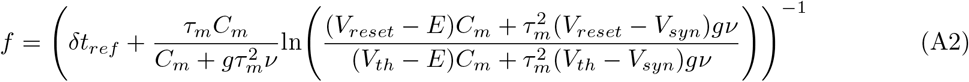

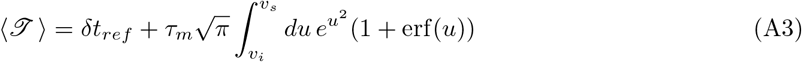

## Appendix B Inhibitory-Excitatory balance

When we study the case of two populations, three different configurations of the network connectivity are analyzed. They are sketched in figure 5. In general, the synaptic dynamics and strength depend both on the population that fires the spike train and the population that receives the stimuli.

**Fig. 5.**
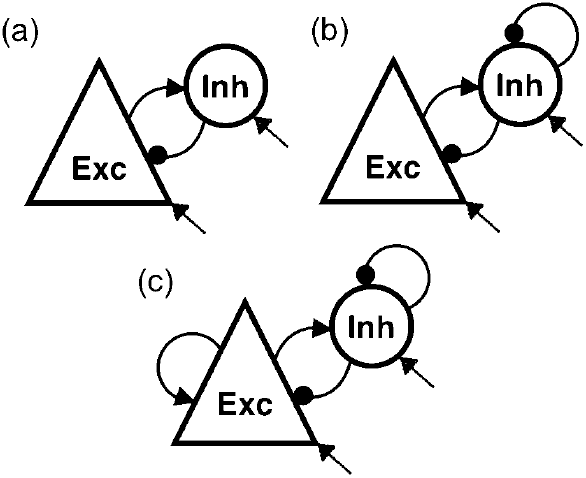
Different types of connectivity. Schematic of the distinct connectivity among the inhibitory neurons and the excitatory neurons we tested. The external arrows represent the external excitation from the thalamus.

### B.1 Exc-Inh loop

Following the main steps as in the article by Brunel and Wang (2003), we introduce a presynaptic firing rate for the inhibitory and excitatory neurons of the form:

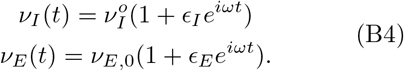

Even when each single neuron fires in a random way, simulations show an oscillatory behavior of the entire network for a certain range of the parameter space.

We require *ω* as the oscillation frequency for both excitatory and inhibitory neurons. As we see by simulations, the two neural classes are substantially synchronized apart from a phase lag. Using equation B4 in equation 6, we then obtain two distinct equations for the variable *s*(*t*) through which the synapses act on neurons:

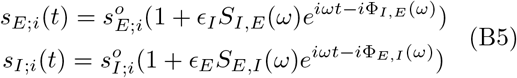

where

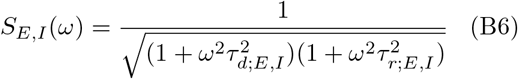

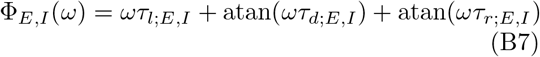

and the synaptic times *τ*_*r*_, *τ*_*d*_ and *τ*_*l*_ have been defined by both presynaptic and postsynaptic populations. The currents of the network then become:

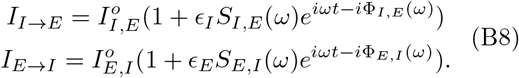

We use only the main oscillatory frequency of the network, and we adopt the approximation of constant membrane potential in the current equation. From now on, we make the current sign explicit adding a phase: zero for the excitatory current, which is positive, and *π* for inhibitory neurons, giving a negative current contribution. Then, the oscillatory components in the excitatory and inhibitory neural activity are related by:

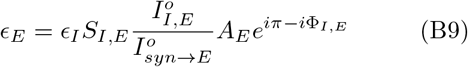

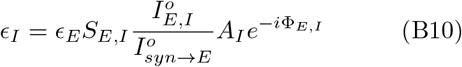

where *A*_*i*_ is a proportional coefficient. Solving these two equations yields the phase condition:

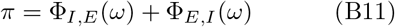

Eq. (B11) is plotted in figure 2 (a) together with the single population phase lag case in black to better compare the two analytical solutions. The difference in population frequency is clearly visible: the population frequency decreases from 180 Hz in the completely inhibitory network (Brunel and Wang, 2003) to 70 Hz, signifying slower activity in the excitatory-inhibitory loop.

### B.2 Adding self-inhibition

We now add recurrent inhibition, and the connectivity becomes as in figure 5(b).

In this system, the synaptic currents are: two inhibitory currents *I*_*I→I*_, *I*_*I→E*_ and one excitatory current *I*_*E→I*_. To the equations in B8 we add:

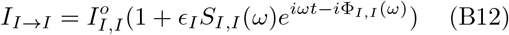

Then, we find:

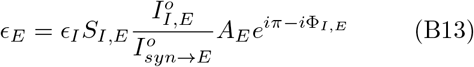

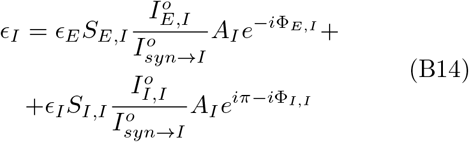

where *A*_*I*_ and *A*_*E*_ are constant coefficients. We define:

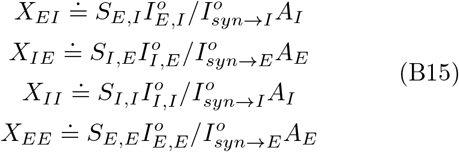

so that using Eq. (B13) into Eq. (B14) we find:

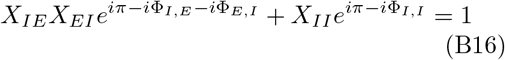

We can solve the equation taking into account its real and imaginary part. When the synapses do not depend on the target population so that Φ_*I,E*_= Φ_*I,I*_, solving in Φ_*I*_, the equations become:

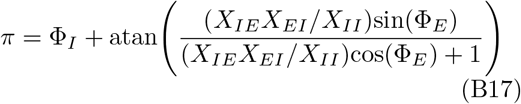

where atan is the arctangent function.

Recurrent inhibition is the reason for the first term on the right side of equations B17, and the excitatory - inhibitory loop produces the other term, which depends on the amplitude B6 of the connections:

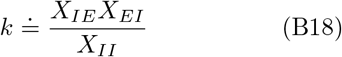

and the phase of the excitatory cells. This contribution can modify the response speed of the network. In figure 2 (a) we display in light-gray the phase-lag curve of Eq. (B17) for a particular value of the connection strength *k* = 2. In figure 2 (b), the population frequency is plotted as a function of *k*. The population frequency decreases for greater values of *k* when loop interactions exceed recurrent inhibition. However, multiple frequencies appear when the strength of the loop interaction is strong enough.

### B.3 Adding self-excitation

Taking into account all the possible connections, we have to include recurrent excitation (see figure 5(c)).

In particular, we add to the previous system the current equation:

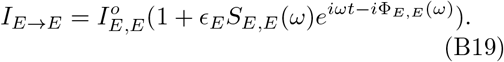

Equaling the presynaptic firing rate to the postsynaptic firing rate, we obtain two equations in *ϵ*_*I,E*_ and the system is self-consistent if:

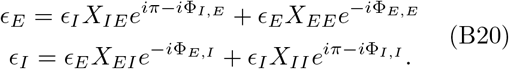

For a certain range of the parameter values the system does not slow down and manifests hyperactivity, signifying epileptic behavior. However, we can reach an equilibrium between the excitation and inhibition currents both on excitatory and inhibitory neurons. Analytically, we can fix the same balance for both neural classes:

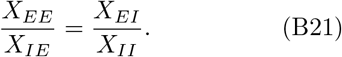

Under this condition, we can simplify the equations B20 to:

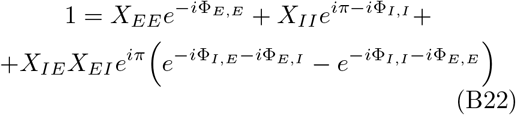

We can further assume that *A*_*E*_ = *A*_*I*_, that is, the network response to the presynaptic stimulus is the same for both neural classes. Additionally, the equation is significantly simplified if synapses do not depend on the target population because the last term in the r.h.s. of the previous equation drops. When the real and imaginary parts are made clear we have:

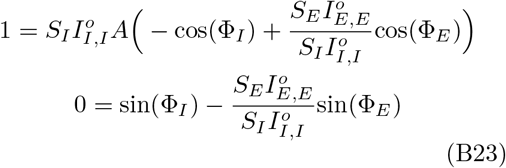

For instance, in the model of Brunel and Wang (2003), the equations B23 are used.

We have studied the solution of equations B23 graphically in figure 2 (c-d). In particular, we want to obtain a value of the frequency of the population activity that satisfies both the real and imaginary parts. However, there are two unknown parameters: the ratio 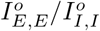 and the amplification factor *A*, which have been estimated by trial and error.

### B.4 Final connectivity

To test our network response, we start with a homogeneous connectivity, 𝒫 = 0.2, and a Poissonian input firing rate of mean 7.3 Hz. The network does not show any kind of remarkable activity, and single neurons have a very low mean firing rate, less than a few Hertz. We then try a batched connectivity depending on the target population, with reference to Billeh et al. (2020). We fix the probability of connection to parvalbumin as 0.4 and to the excitatory neurons as 0.2. If we increase the input firing rate to 30 Hz, the network shows a more pronounced activity, but the mean firing rate of single neurons is still very low. Moreover, the power spectrum shows the most pronounced frequencies at approximately 60 Hz.

We then study the network behavior according to some parameter changes. We introduce a scaling factor both on the connection probability and on the conductance *g* to homogeneously vary the connectivity features, see Fig. S8 in the supplementary material (Online Resource). The external mean firing rate is also shifted from a few Hertz to almost half a hundred Hertz to study the response to external stimulus variations. Starting from batched connectivity, we refine the connection probabilities to the Exc neurons, and finally, the connection probability to the Pvalb neurons is slightly modified. Different parameters are tested: 𝒫_*P,P*_, 𝒫_*P,E*_ and 𝒫_*E,E*_. The fourth 𝒫_*E,P*_ does not show particular phenomenona. While the main frequency of population activity slightly evolves due to the parameter changes, except in a few cases, the single neuron mean firing rate strongly varies. Decreasing the strength of the connections or their probability makes single neurons more active, while synchrony decreases. In fact, the recurrent inhibition decreases.

Additionally, the external input has an important role since it can cause a sharp phase transition in single neuron activity. If we intensify recurrent inhibition of Pvalb or recurrent excitation on Exc it is likely to expect for an increase in single cell activity. It happens sharply in the present network.

## Appendix C Build the three populations network

Figure 6 depicts the connectivity we used to study the case of three populations.

**Fig. 6.**
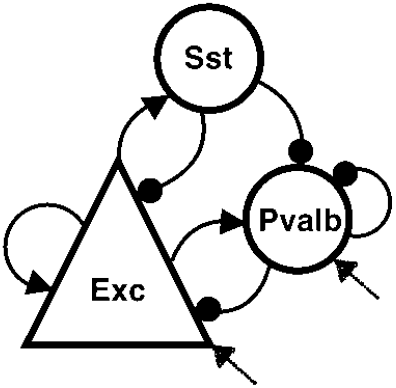
Three populations connectivity. The new model is supposed to connect almost every type of neuron, expect for the recurrent inhibition of somatostatin neurons and the inhibition of somatostatin by parvalbumin neurons. Moreover from the literature, the external input does not affect the somatostatin neurons. The external arrows show the excitation from the thalamus.

Somatostatin cells are stimulated by the excitatory neurons, and their firing influences in turn the spiking activity of both parvalbumin and excitatory neurons; the excitatory - parvalmbumin subnetwork is completely interconnected. The other connection probabilities are almost one order lower (Billeh et al., 2020) and they are set equal to zero: 𝒫_*P,S*_ = 𝒫_*S,S*_ = 0.

To determine the actual connection probability, a recursive study is performed.

1. Starting from a connection probability among Exc and Pvalb as in the case of two populations, we add the excitation of Sst by Exc and analyze their response to different stimuli. In particular, both the conductivity and the probability of connection 𝒫_*E,P*_ are varied.
2. The next step is redrawing the feedback from Sst to Pvalb and from SSt to Exc. This is first done separately for the two target populations. When not varied, *𝒫*_*S,P*_ = 0 and *𝒫*_*S,E*_ = 0.
3. We now impose 𝒫_*S,P*_ = 0.4 and 𝒫_*S,E*_ = 0.2, fixing the two connection probabilities at the same time. Then, a scaling factor is included, and we modify both the connection probability and the conductance.
4. Finally, the connection probability is fixed equal to *𝒫*_*S,E*_ = 0.15, *𝒫*_*S,P*_ = 0.03.

There are other parameter values we can changed, first, the conductivity from Sst to Pvalb neurons, but they do not show particular behaviors.

We can also modify the connection probability among Pvalb and Exc neurons, yet we have decided to use the connectivity of the final network in the case of two populations (Pvalb - Exc) as we have built it in the previous section, and add a new contribution from the Somatostatin class, instead of defining again the entire connectivity. See the the supplementary material (Online Resource).

### C.1 Analytical approach

The three synaptic currents for the present network are written below, the usual format being used (we neglect the fluctuation terms):

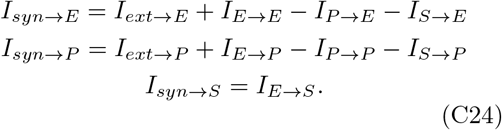

As in the previous sections, currents are supposed to be linearly dependent on the input firing rate, that we assumed to be of the form:

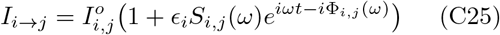

and the network response to the current stimulus is supposed to be linear.

The postsynaptic firing rate is obtained:

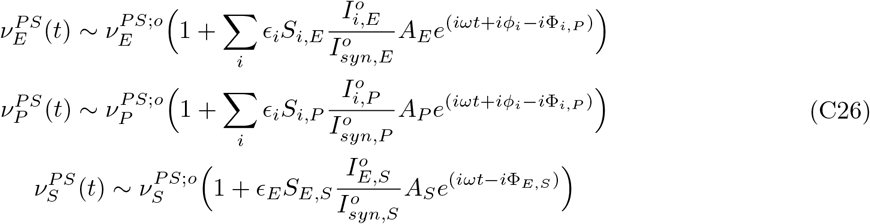

where the index runs over the three populations and the phase lag *ϕ* is zero when the presynaptic population is excitatory and *π* otherwise for inhibitory presynaptic populations. From the three postsynaptic firing rates, we automatically derive the phase lag condition. In particular, the system of equations below has to be resolved:

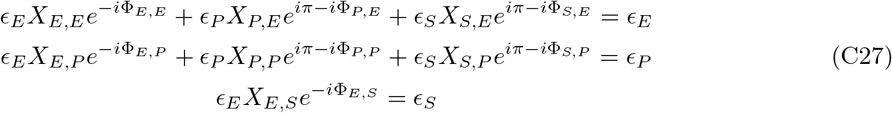

and then we find:

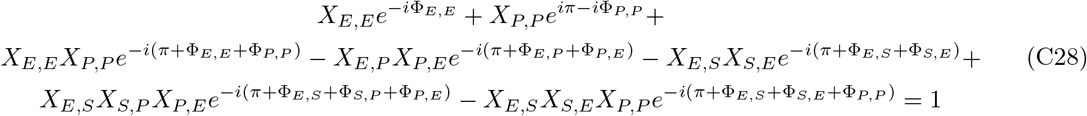

