## Supplementary-Material for "A spiking LIF model captures the role of Somatostatin and Parvalbumin neurons in generating oscillations in V1"

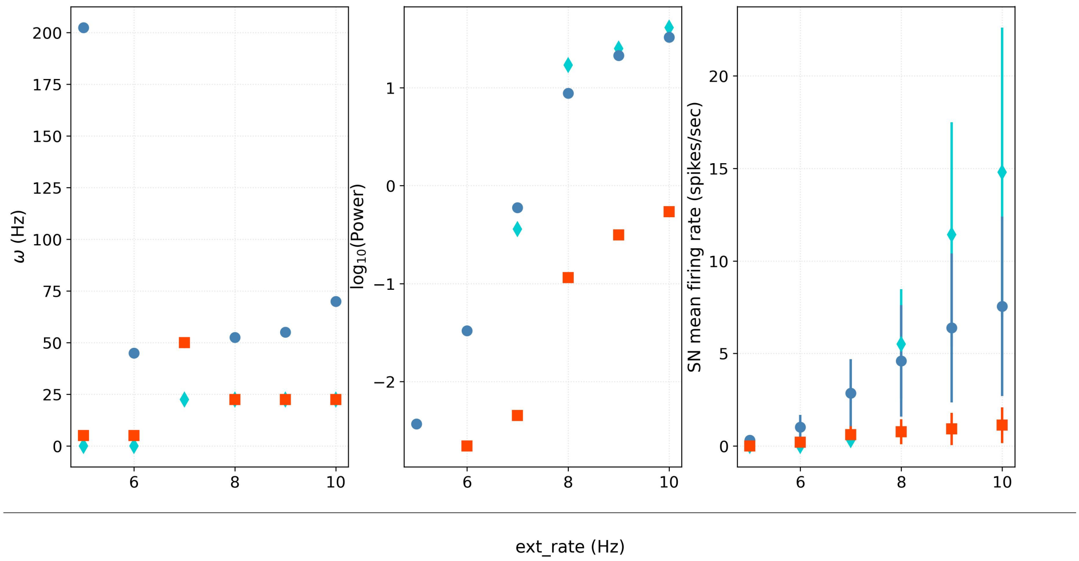


**Fig. S 1 Three-populations analysis**
Network behavior is studied for different values of the input mean external rate. The external stimulation is a Poissonian distribution of excitatory spikes whose mean rate is constant over time. The mean external rate goes from 5 Hz to 10 Hz. Network connectivity is the same as in Figure 4 of the main text. In blue dots we show the parvalumin neurons, in cyan diamonds the somatostatin neurons and in red squares the excitatory neurons. In the left part of the figure we show the peak in the oscillation frequency: it rapidly stabilizes at approximately 50 Hz for paravalbumin neurons and 25 Hz for somatostatin and excitatory neurons. The central figure represents the power related to $\omega$. Note that for low input rate the power is two orders of magnitude less than for high input rate. On the right side it is shown the single neuron mean firing rate (the bars represents the $75^{th}$ percentile). It increases almost linearly with the input rate.

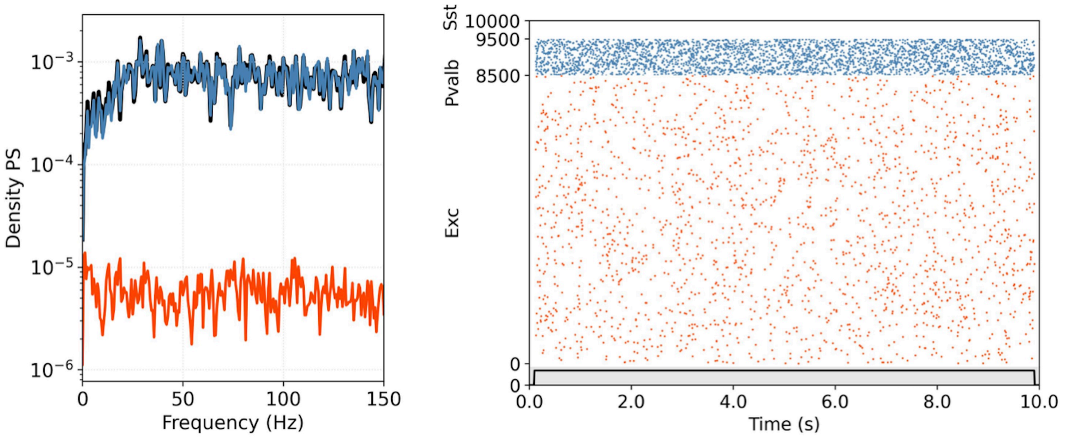


**Fig. S 2 Three-populations 5 Hz input rate**
The raster plot and its power spectrum are shown when the external input rate is set equal to 5 Hz. Network connectivity is the same as in Figure 4 of the main text. In blue we show the parvalumin neurons, in cyan the somatostatin neurons and in red the excitatory neurons. Note how there is no contribution of the somatostatin neurons and also parvalbumin and excitatory neurons do not show any particular oscillatory behavior.

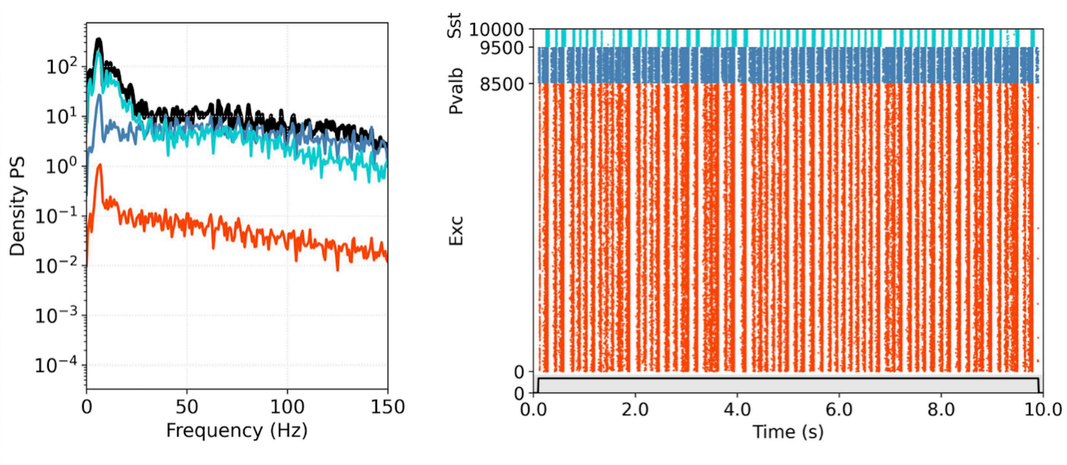


**Fig. S 3 Three-populations 10 Hz input rate**
The raster plot and its power spectrum are shown when the external input rate is set equal to 10 Hz. Network connectivity is the same as in Figure 4 of the main text. In blue we show the parvalumin neurons, in cyan the somatostatin neurons and in red the excitatory neurons. Somatostatin neurons contribute significantly to network oscillations with frequencies lower than approximately 25 Hz.

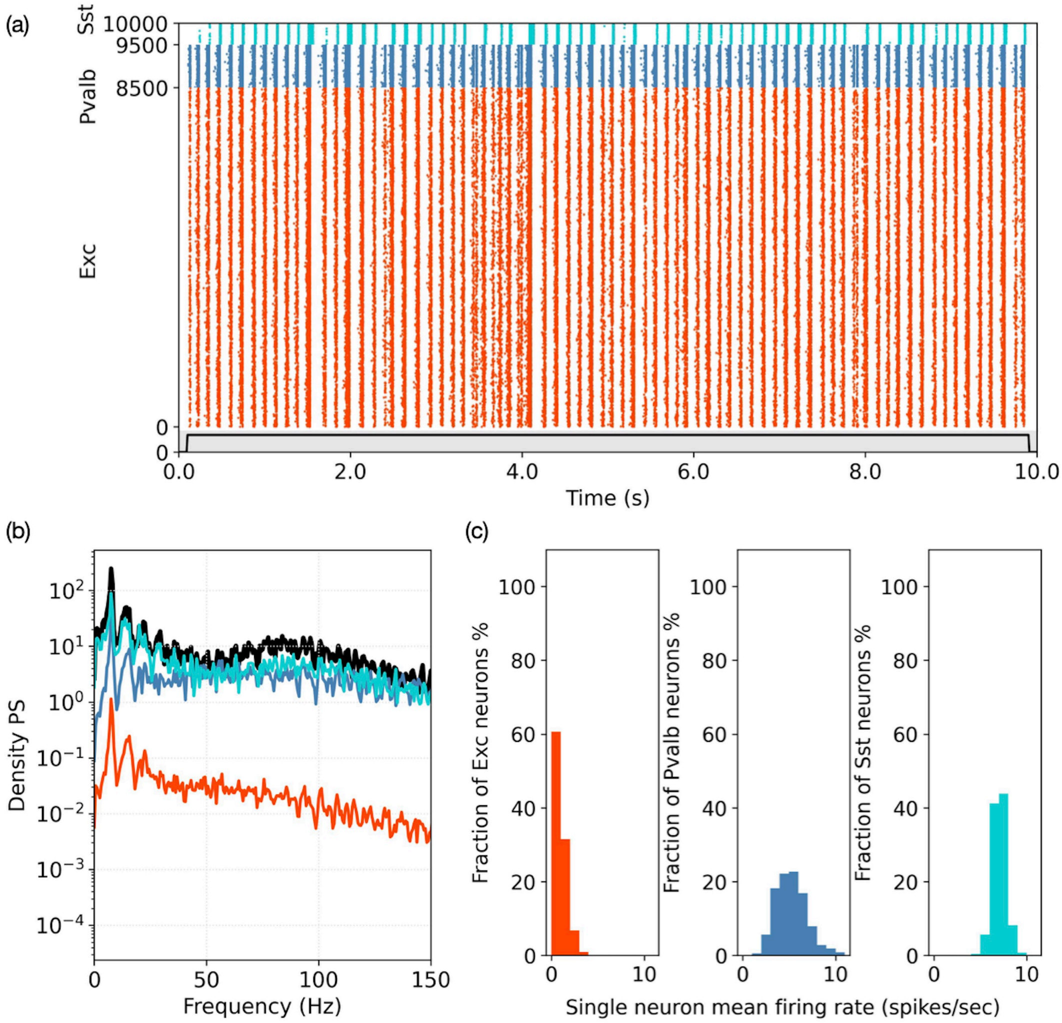


**Fig. S 4 Hyperpolarized Pvalb neurons with 100 pA current**
The connection probability is the same as state in Figure 4 in the main text. The mean rate of the external excitatory input is set equal to 7.3 Hz. An external constant current stimulus of amplitude equal to 100 pA is added to the Pvalb neurons.
(a) Raster plot of the entire network made of 1000 parvalbumin neurons, 500 somatostatin neurons and 8500 excitatory neurons.
(b) Power spectrum (Welch's method applied) of the population activity. Red refers to the excitatory neurons, blue to the parvalbumin neurons and cyan to the somatostatin neurons. The black line shows the oscillation components of the entire network response.
(c) Histograms of the single neuron mean firing rate for excitatory, parvalbumin and somatostatin neurons.

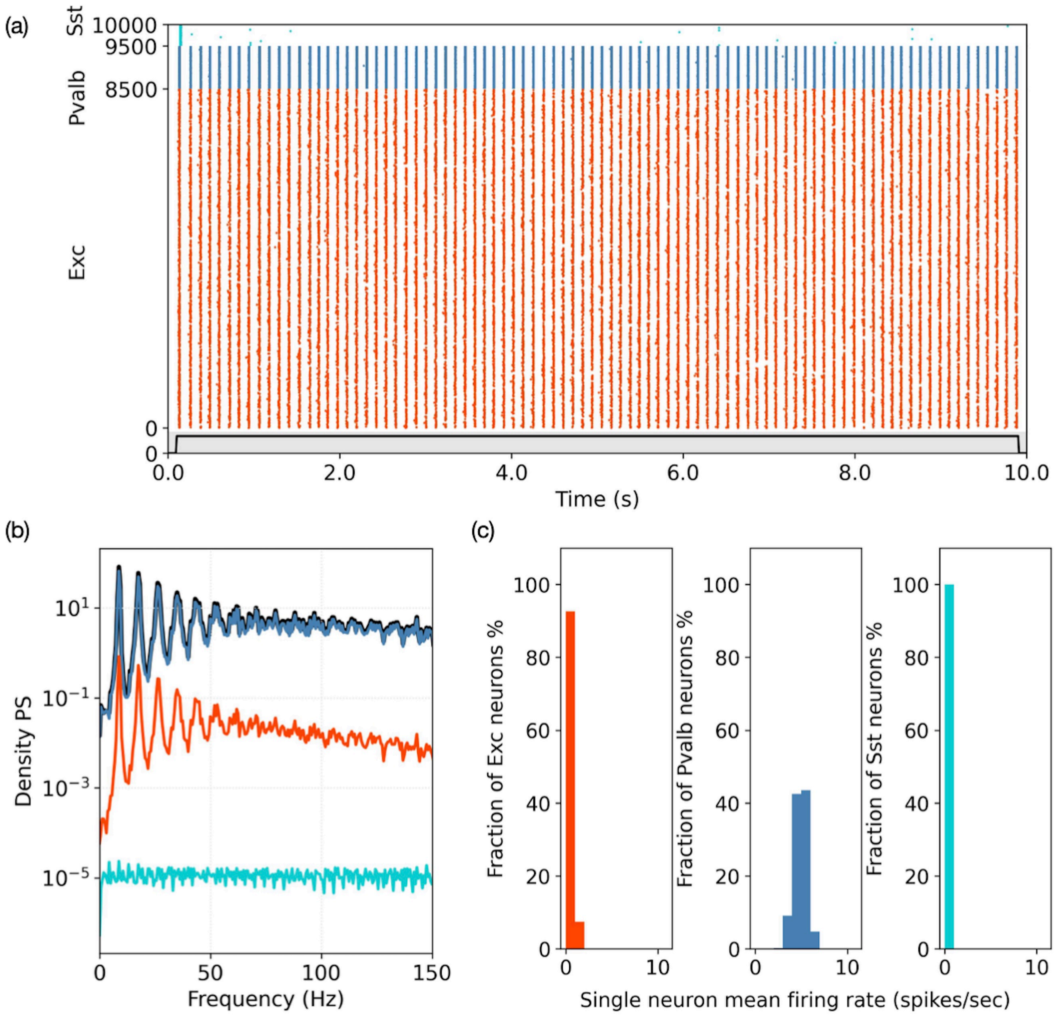


**Fig. S 5 Hyperpolarized Pvalb neurons with 200 pA current**
The connection probability is the same as state in Figure 4 in the main text. The mean rate of the external excitatory input is set equal to 7.3 Hz. An external constant current stimulus of amplitude equal to 200 pA is added to the Pvalb neurons.
(a) Raster plot of the entire network made of 1000 parvalbumin neurons, 500 somatostatin neurons and 8500 excitatory neurons.
(b) Power spectrum (Welch's method applied) of the population activity. Red refers to the excitatory neurons, blue to the parvalbumin neurons and cyan to the somatostatin neurons. The black line shows the oscillation components of the entire network response.
(c) Histograms of the single neuron mean firing rate for excitatory, parvalbumin and somatostatin neurons.

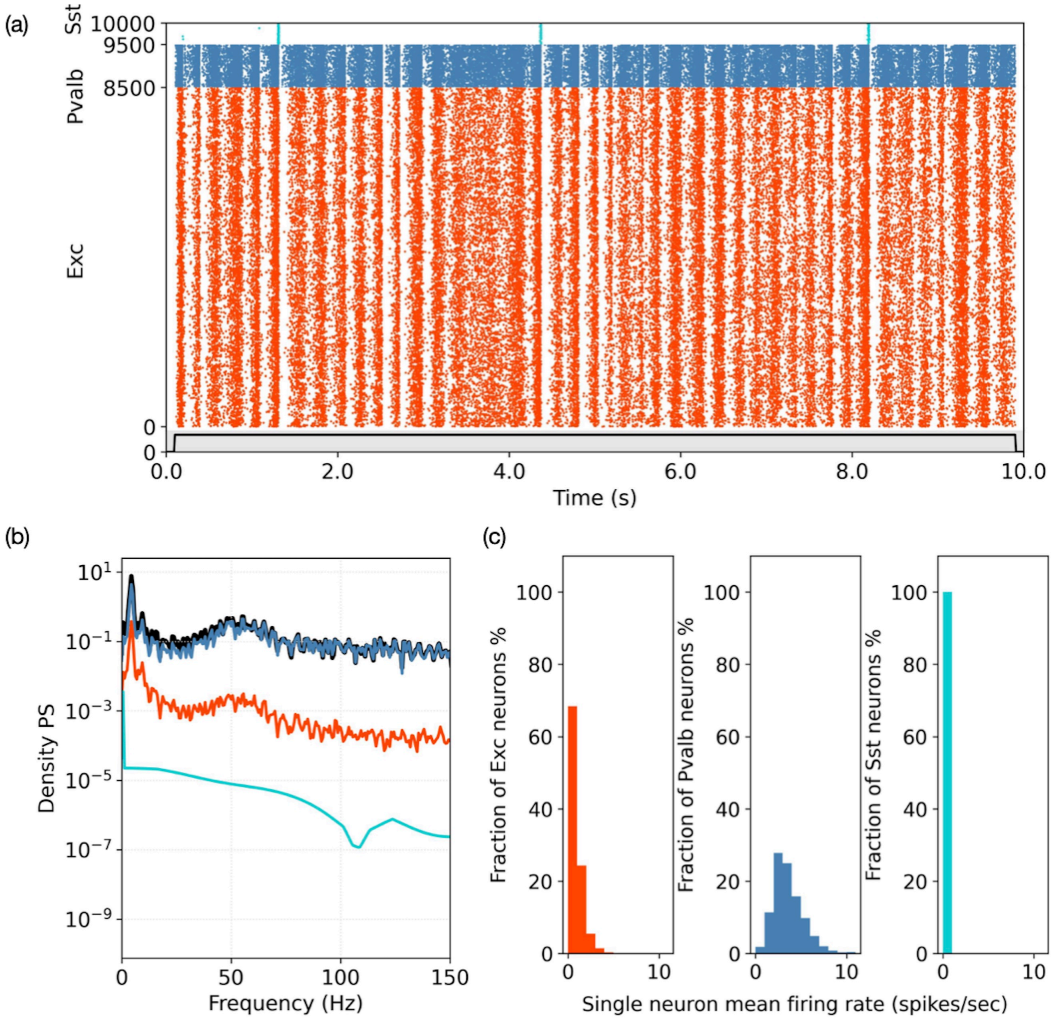


**Fig. S 6 Hyperpolarized Sst neurons with 100 pA current**

The connection probability is the same as state in Figure 4 in the main text. The mean rate of the external excitatory input is set equal to 7.3 Hz. An external constant current stimulus of amplitude equal to 100 pA is added to the Sst neurons.

(a) Raster plot of the entire network made of 1000 parvalbumin neurons, 500 somatostatin neurons and 8500 excitatory neurons.

(b) Power spectrum (Welch's method applied) of the population activity. Red refers to the excitatory neurons, blue to the parvalbumin neurons and cyan to the somatostatin neurons. The black line shows the oscillation components of the entire network response.

(c) Histograms of the single neuron mean firing rate for excitatory, parvalbumin and somatostatin neurons.

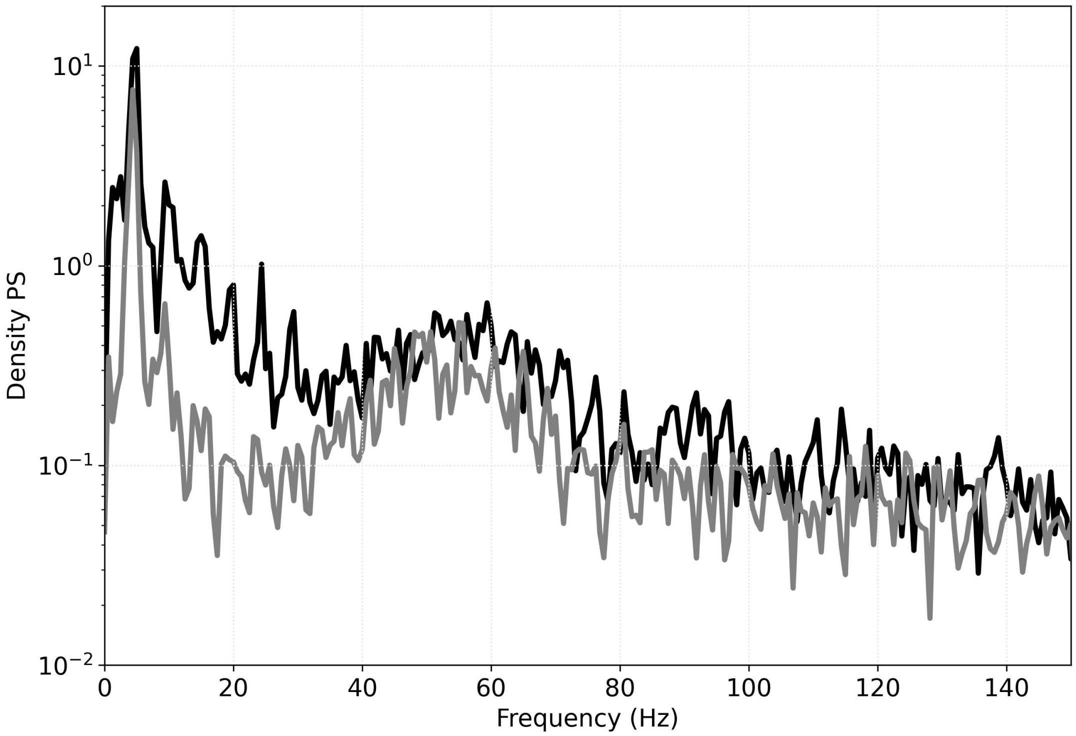


**Fig. S 7 Decrease in beta activity due to hyperpolarization on somatostatin neurons**
The black curve represents the power spectrum in the standard setup of the three-populations network. The gray curve is the power spectrum when somatostatin neurons are hyperpolarized by a constant current equals to 100 pA. Both cases count all the contributions due to parvalbumin, somatostatin and excitatory neurons. Note how gamma frequencies are almost not affected while frequencies in the beta range decrease significantly.

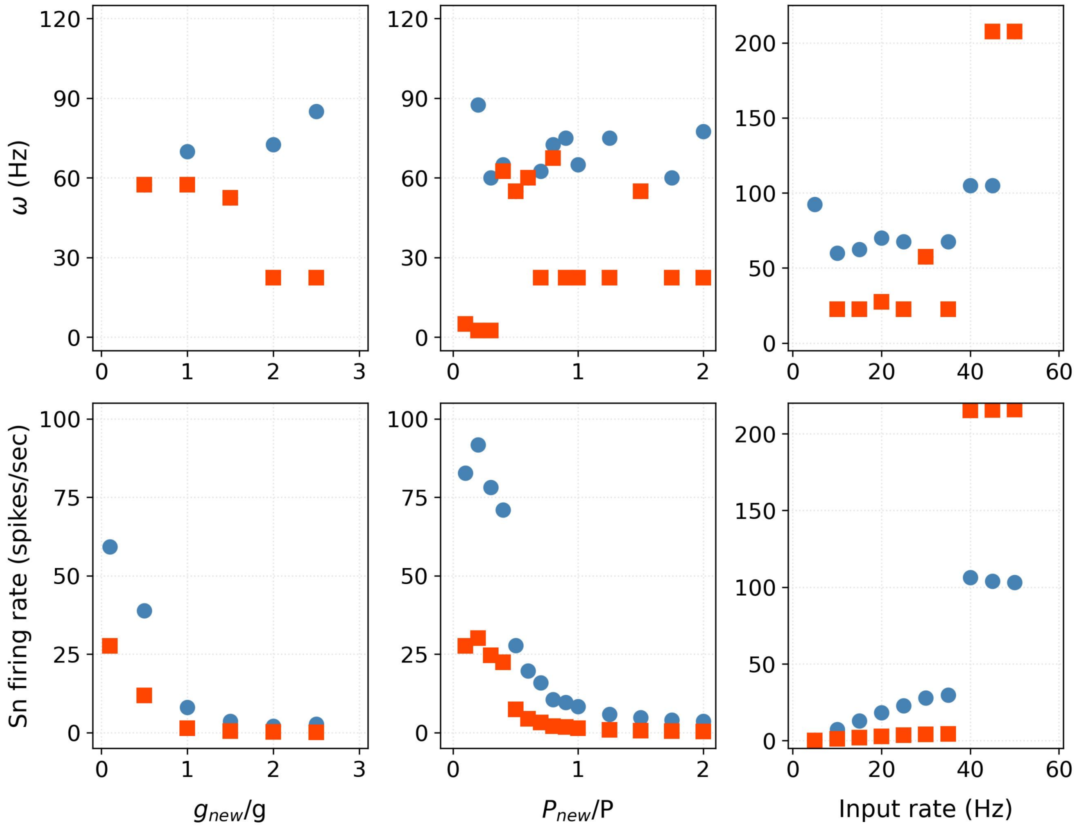


**Fig. S 8 Two-populations systematic study**
The network behavior is studied according to some parameter changes: the conductance g, the connection probability P and the external input rate. Single neuron and population activity are shown: in blue dots the Parvalumin and in red squares the Excitatory neurons. In particular note how the increase of external input rate at approximately 40 Hz determines a fast increase in the single neuron mean firing rate.

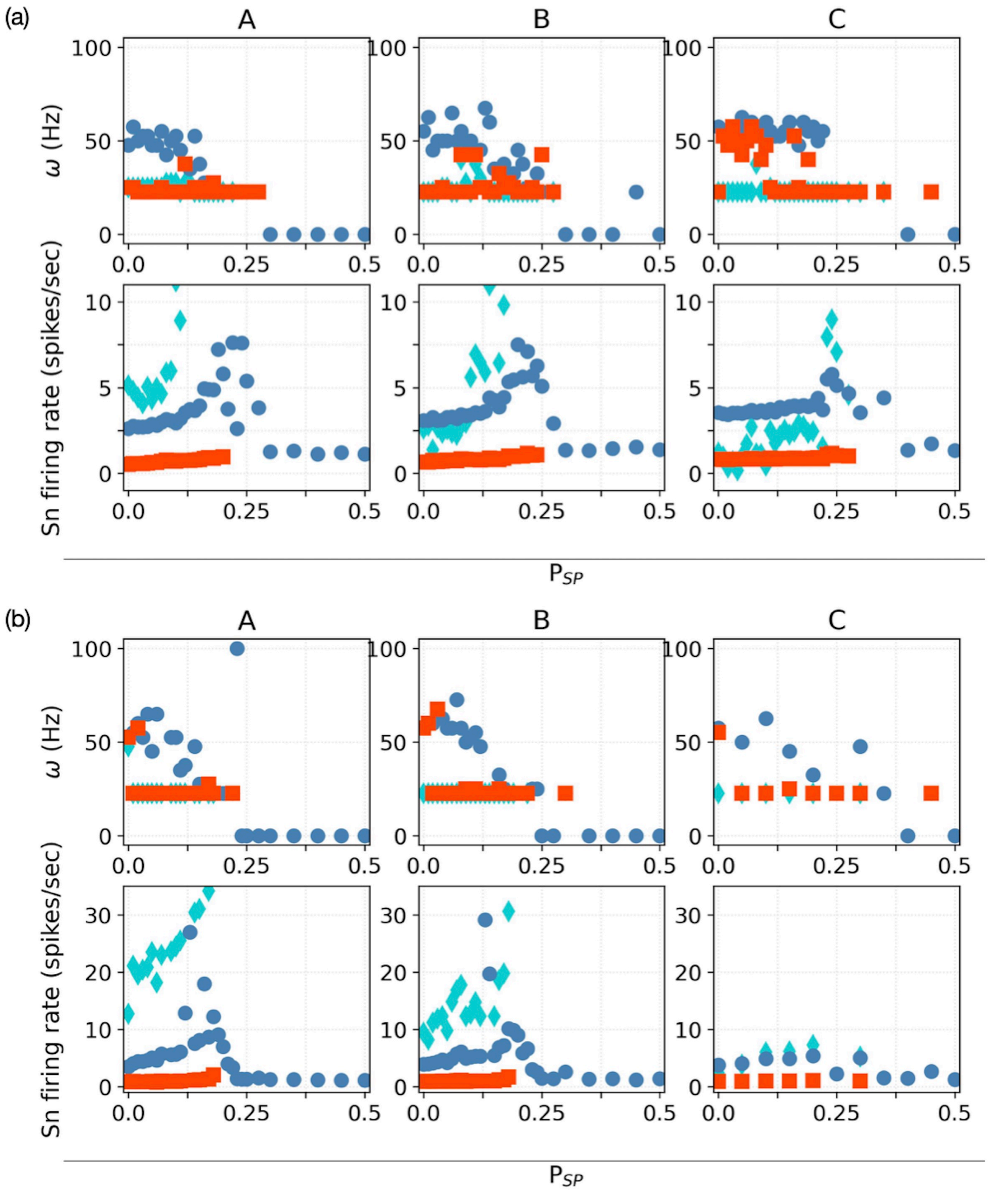


**Fig. S 9 Three-populations systematic study**
Single neuron and population activity for distinct sets of parameter values are shown. In blue dots are the Pvalb neurons, in cyan diamonds the Sst neurons and in red squares the Exc neurons. Columns stand for: A $\mathcal{P}_{E,S} = 0.24$ , B $\mathcal{P}_{E,S} = 0.18$, C $\mathcal{P}_{E,S} = 0.12$. In (a) $\mathcal{P}_{S,E} = 0$. In (b) $\mathcal{P}_{S,E} = 0.05$.

(a)

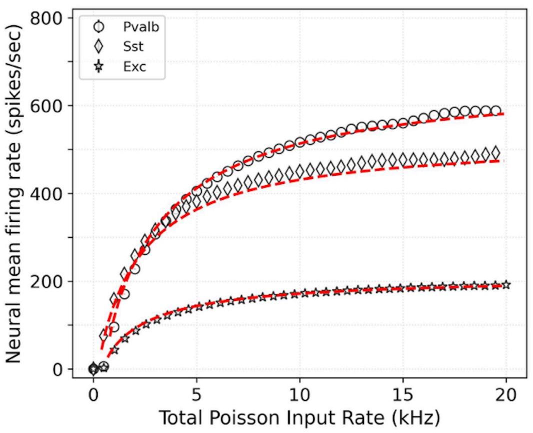


(b)

| | Pvalb | Sst | Exc |
|---|---|---|---|
| reference Tau (ms) | 7.85 | 16.82 | 21.49 |
| estimated Tau (ms) | 8.5 | 15.0 | 24.6 |
| +/- | 0.1 | 0.7 | 0.2 |

(c)

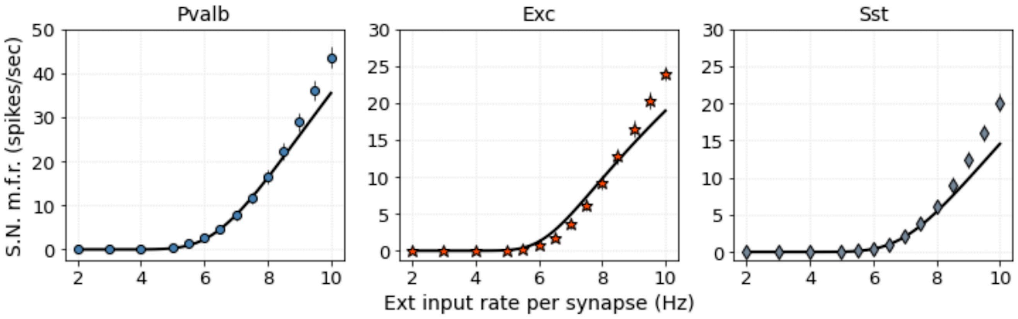


**Fig. S 10 Single neuron analysis**
(a) The markers identify the neural mean firing rate for different values of the total mean input firing rate - we take into account the number of the external synapses -. The dashed lines shows the results of fitting using the equation $f = \left(\delta t_{ref} + \frac{\tau_m C_m}{C_m + g\tau_m^2 \nu} \ln\left(\frac{(V_{reset}-E)C_m + \tau_m^2 (V_{reset}-V_{syn})g\nu}{(V_{th}-E)C_m + \tau_m^2 (V_{th}-V_{syn})g\nu}\right)\right)^{-1}$, where $\tau_m$ is set as free parameter. The analytical approximation fits relatively well the range of strong input as we can see from the results of the fit. (b) The first row shows the reference value of characteristic membrane time $\tau_m$ for every neural type as we have deduced from literature. The second and third rows display the estimated values with error using the curve fit procedure in python. (c) The single neuron mean firing rate is shown for different sets of external input rate per synapse. The other parameters are as usual. To describe the response of Somatostatin neuron to the external stimulus we use the same synaptic type as for the Parvalbumin neuron. The markers show the mean and standard deviation calculated for 1000 trials. The black curves correspond to the analytical solution of the expected frequency $\nu_0 = \langle \mathscr{T} \rangle^{-1}$, obtained by the Fokker-Planck model through equation: $\langle \mathscr{T} \rangle = \delta t_{ref} + \tau_m \sqrt{\pi} \int_{v_i}^{v_s} du\, e^{u^2}(1 + \mathrm{erf}(u))$.
